# Genomic and phenotypic diversification of *Pseudomonas aeruginosa* during sustained exposure to a ciliate predator

**DOI:** 10.64898/2026.01.13.699197

**Authors:** Nayma Romo Bechara, Nikhil Bardeskar, Heather A. Hopkins, Louis-Marie Bobay, Kasie Raymann

## Abstract

Predator-mediated selection is an important ecological force shaping bacterial evolution, but its effects on genomic adaptation and virulence in opportunistic pathogens are not fully understood. Here, we used experimental evolution to study how exposure to the ciliate predator *Tetrahymena thermophila* affects *Pseudomonas aeruginosa*. Replicate populations were evolved for 60 days with or without the predator, followed by whole-genome shotgun metagenomic sequencing and phenotypic analyses.

Both treatments showed strong selection and evidence of parallel evolution at gene and nucleotide levels, indicating constrained adaptation. However, predator exposure altered evolutionary dynamics. Predator-evolved populations showed a wider distribution of mutation frequencies, with many mutations persisting at intermediate frequencies, consistent with increased clonal interference and ongoing competition among lineages. In contrast, populations evolved without predators showed more high-frequency mutations, consistent with selective sweeps, though some low-frequency variants remained.

Despite substantial genomic change, phenotypic outcomes were variable. Virulence in an invertebrate host model did not consistently increase; instead, evolved isolates showed context-dependent changes, including modest decreases or occasional increases. Competition assays also showed no consistent fitness advantage for predator-evolved isolates, suggesting trade-offs between predator resistance and growth in other environments.

Overall, predator-mediated selection reshaped evolutionary dynamics by maintaining diversity and altering the balance of lineages rather than producing uniform increases in virulence. These results highlight how ecological complexity influences adaptive evolution and the context-dependent nature of pathogen traits.

**Importance:** Opportunistic pathogens like *Pseudomonas aeruginosa* often evolve in environmental settings before infecting hosts, raising questions about how ecological interactions influence virulence. Predator-mediated selection has been suggested to increase virulence via coincidental evolution, but evidence is inconsistent. Here, we show that exposure to a eukaryotic predator does not consistently elevate virulence but does reshape evolutionary dynamics by altering how mutations spread in populations. Predator-exposed populations retained more intermediate-frequency mutations, consistent with increased clonal interference and ongoing competition among lineages, whereas non-predator populations were dominated by selective sweeps. These differences were also reflected in functional targets of adaptation, with predator exposure favoring mutations in genes involved in environmental sensing and interaction. Together, these findings suggest that ecological complexity shapes the dynamics of adaptation rather than driving a single evolutionary outcome, highlighting that virulence is an emergent property influenced by underlying evolutionary processes.

## Introduction

*Pseudomonas aeruginosa* is a ubiquitous and highly versatile opportunistic pathogen that inhabits a wide range of environmental and host-associated niches, including soil, aquatic systems, plant surfaces, and animals (Silby et al. 2011; Wu and Li 2015; Crone et al. 2020; Diggle and Whiteley 2020). Its ecological success is largely driven by a large and flexible genome encoding diverse metabolic capabilities, regulatory networks, and virulence factors, allowing rapid adaptation to changing environments (Stover et al. 2000; Coggan and Wolfgang 2012; Aroca Molina et al. 2024). In clinical settings, *P. aeruginosa* is a major cause of opportunistic and nosocomial infections and a leading contributor to morbidity and mortality in individuals with cystic fibrosis (Kerr and Snelling 2009). Understanding how environmental selection shapes the evolution of this organism is therefore critical for linking ecological processes to pathogenic potential.

Experimental evolution studies have demonstrated that *P. aeruginosa* adapts rapidly to diverse selective pressures, including nutrient limitation (Cecil et al. 2025), antibiotic exposure (Barbosa et al. 2017), oxidative stress (Chua et al. 2016; Fu et al. 2024), and host-associated environments (Marvig et al. 2014). These studies have revealed predictable genetic changes, particularly in regulatory pathways, and have highlighted the importance of environmental context in shaping both genomic and phenotypic evolution. Among these selective pressures, interactions with microbial predators such as protozoa, nematodes, and bacteriophages have been increasingly recognized as key drivers of bacterial diversification (Matz and Kjelleberg 2005; Mikonranta et al. 2012; Williams 2013). Predator-mediated selection can influence traits closely linked to pathogenicity, including motility, biofilm formation, toxin production, and surface modification (Matz et al. 2004; Jousset et al. 2009), and has been proposed as a major ecological force structuring microbial communities (Nair et al. 2019).

These observations form the basis of the coincidental evolution hypothesis, which posits that virulence in opportunistic pathogens may arise as a by-product of traits selected in environmental contexts, including defense against eukaryotic predators (Levin 1996; Hosseinidoust et al. 2013; Brown et al. 2007). Under this framework, certain adaptations that enhance survival during predation, such as resistance to phagocytosis, secretion of toxins, or biofilm formation, may incidentally increase the ability of bacteria to colonize or damage animal hosts. However, empirical support for this hypothesis remains mixed. While some studies have shown that predator exposure can enhance or maintain virulence-associated traits (Amaro et al. 2015; Nair et al. 2019; Hoque et al. 2021; Hopkins et al. 2024), others have reported reduced or context-dependent effects (Mikonranta et al. 2012; Friman and Buckling 2014; Granato et al. 2018; Leong et al. 2022; Saha et al. 2025), suggesting that the relationship between environmental selection and virulence is more complex than originally proposed.

One potential explanation for these contrasting outcomes is that predator-mediated selection influences not only which traits evolve but also how evolution proceeds. Theoretical and empirical work in experimental evolution indicates that simple, stable environments often promote rapid selective sweeps and repeatable outcomes, whereas more complex environments, particularly those involving biotic interactions, can maintain multiple competing lineages through clonal interference and expand the range of accessible adaptive solutions (Gerrish and Lenski 1998; Desai and Fisher 2007; Good et al. 2017). In such systems, adaptation may generate greater genetic and phenotypic diversity, leading to variable rather than directional changes in traits such as virulence.

Despite growing interest in predator-driven evolution, relatively few studies have examined how sustained exposure to eukaryotic predators shapes both the genomic and phenotypic evolution of *P. aeruginosa*, particularly at the population level. Existing work with amoebae (*Acanthamoeba castellanii*) and nematodes (*Caenorhabditis elegans*) has demonstrated that predator or host interactions can alter virulence-associated traits (Matz et al. 2008; Granato et al. 2018; Leong et al. 2022), but the evolutionary consequences of long-term interaction with generalist protist predators remain poorly understood. In particular, it is unclear whether predator-mediated selection drives consistent increases in virulence, maintains phenotypic diversity, or interacts with other selective pressures such as nutrient adaptation.

Here, we address this gap using experimental evolution to investigate how exposure to the ciliate predator *Tetrahymena thermophila* shapes the evolution of *P. aeruginosa*. Populations were evolved for 60 days (~276 generations) in the presence or absence of the predator, and whole-genome shotgun (WGS) metagenomic sequencing and phenotypic assays were used to characterize genomic changes and their effects on fitness and virulence-related traits. Our results show that predator presence does not simply alter the targets of selection, rather reshapes the dynamics of adaptation. Predator-exposed populations exhibited a broader distribution of mutation frequencies, with a greater proportion of mutations maintained at intermediate frequencies, consistent with increased clonal interference and sustained competition among lineages. By contrast, populations evolved without the predator displayed a more pronounced bimodal distribution, with enrichment of high-frequency mutations indicative of selective sweeps while retaining a subset of lower-frequency variants. Despite these differences, adaptation remained strongly constrained at the genetic level, with repeated targeting of key regulatory and virulence-associated genes. Importantly, changes in virulence were modest and context dependent, with both increases and decreases observed across isolates and host conditions. Together, these findings suggest that predator-mediated selection does not uniformly drive the evolution of virulence, rather alters the dynamics of adaptation, generating diverse evolutionary trajectories that can produce variable host-associated outcomes. By linking evolutionary dynamics, genomic adaptation, and phenotypic expression, this study provides new insight into how ecological complexity shapes the emergence and maintenance of virulence in opportunistic pathogens.

## Results

### Experimental evolution and genome sequencing

Six media-evolved (ME) and six predator-evolved (PE) replicate populations of *P. aeruginosa* strain NRB were maintained by daily passaging. In PE lines, 4% of each bacterial culture was transferred into fresh axenic *T. thermophila* cultures each day to minimize predator–prey co-evolution. Successful bacterial transfer and predator viability were verified prior to each passage. All six ME and six PE populations were subjected to WGS metagenomic sequencing after 60 days of evolution (~276 generations).

### Mutation identification and filtering

The ancestral strain was initially sequenced using a combination of long-read (Oxford Nanopore) and short-read (Illumina) technologies. Because the strain was presumed to be *Pseudomonas aeruginosa* PAO1, sequencing reads were first aligned to the PAO1 reference genome; however, alignment was poor, suggesting that the strain was not PAO1 or exhibited substantial genomic divergence. To generate an appropriate reference genome, the sequencing data were assembled de novo using a hybrid assembly approach implemented in hybridSPAdes, integrating both long- and short-read data to produce a high-quality genome assembly. The resulting genome exhibited 99.996% nucleotide identity to *P. aeruginosa* FDAARGOS_1041, differing by approximately 274 nucleotides. Mutations in evolved populations were subsequently identified by aligning sequencing reads to this de novo assembled ancestral reference genome. Variants detected in the ancestral strain during read mapping were excluded to minimize false-positive mutation calls.

To assess potential cross-contamination among populations, we performed clonal deconvolution followed by phylogenetic analysis (see Methods). One ME population (ME6) showed evidence of contamination and was excluded from all subsequent analyses (Figure S1).

### Genome-wide mutation landscape across all detected variants

Using a frequency threshold of ≥0.05 and base quality ≥30, we identified 647 distinct mutations, excluding same-site parallel mutations. The number of mutations per population ranged from 10 to 98, with mean mutation frequencies ranging from 0.10 to 0.79 and an average genome coverage of 94× (Dataset S1; Table S1). The average number of mutations per generation was 0.19 for ME populations and 0.24 for PE populations (Table S2).

The total number of mutations per population was not correlated with average genome coverage (R^2^ = 0.1094, P = 0.32; Figure 1A), suggesting that sequencing depth was sufficient to capture the mutations present in each population. The number of mutations per population was significantly negatively correlated with average mutation frequency (R^2^ = 0.6752, P = 0.0019; Figure 1B). Populations harboring a greater number of mutations tended to maintain these at lower individual frequencies, a pattern consistent with increased genetic diversity and/or ongoing clonal interference, where multiple competing lineages prevent individual mutations from reaching high frequency. By contrast, populations with fewer mutations at higher frequencies may reflect recent selective sweeps, in which one or a small number of advantageous mutations rapidly increased in frequency, reducing overall diversity.

**Figure 1.**
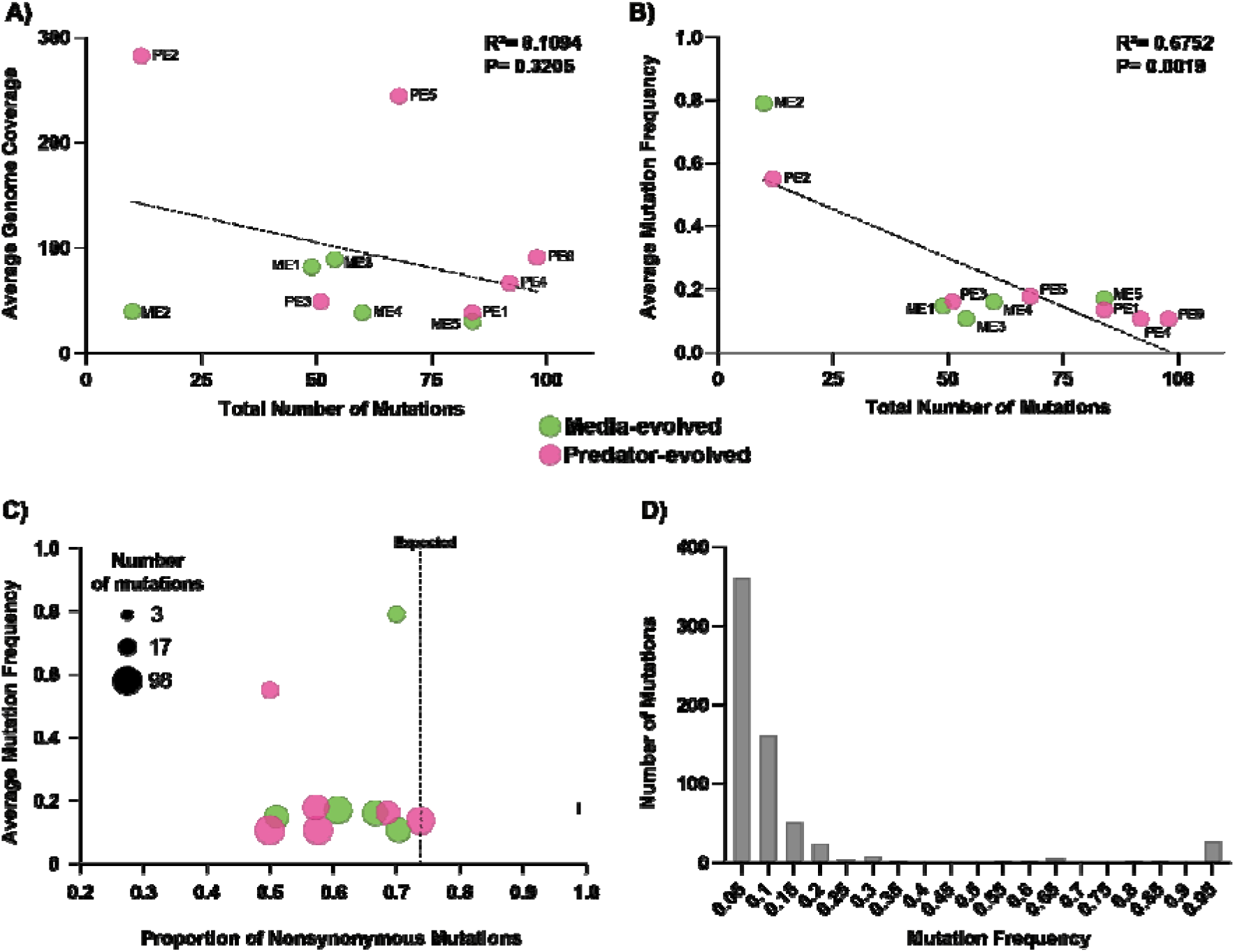
Mutations (≥0.5 frequency) detected in evolved populations. **A)** Relationship between average mutation frequency and the total number of mutations per evolved population. **B)** Relationship between average genome coverage and the total number of mutations per evolved population. For panels A and B, R^2^ values and P-values were calculated using simple linear regression. **C)** Mutation frequency and proportion of nonsynonymous mutations across evolved populations. Scatter plots show the average mutation frequency (y-axis) versus the proportion of nonsynonymous and nonsense mutations (x-axis) for each evolved population at D60. Circle size represents the total number of mutations detected per population. The vertical dashed line represents the number of nonsynonymous (including nonsense) mutations expected by random chance. **D)** Number of mutations detected at different frequencies between 0.5 and 1 across all evolved populations. Each bar represents the lower bound of each 0.05 interval (e.g., 0.05 = 0.05–0.09; 0.95 = 0.95– 1.0).

Mutations in each evolved population were classified as nonsynonymous (including nonsense mutations), synonymous, RNA, or intergenic (Figure 1C; Dataset S1). To estimate the expected proportions of mutations affecting these sequences under random mutation, we performed simulations on the ancestral genome based on the mutation patterns observed in each line. Briefly, for each line, we estimated the observed number of mutations as well as the relative mutability of each nucleotide and the mutation spectrum. These estimates were then used to perform 10,000 simulations for each line, thereby reflecting the mutation biases recorded in our experiments. The proportion of nonsynonymous mutations in the evolved lines ranged from 0.50 to 0.74, compared to an expected 0.74 under random mutation (Figure 1C).

Considering all detected mutations, the frequency distribution was strongly skewed toward low-frequency variants, with the majority of mutations present at frequencies below 0.20 (Figure 1D). Because low-frequency variants may include false positives arising from sequencing noise, coverage variability, or read mismapping and may also represent transient polymorphisms rather than stable mutations, subsequent analyses focused on mutations present at ≥0.20 frequency in at least one population (Figures 2 and 3).

**Figure 2.**
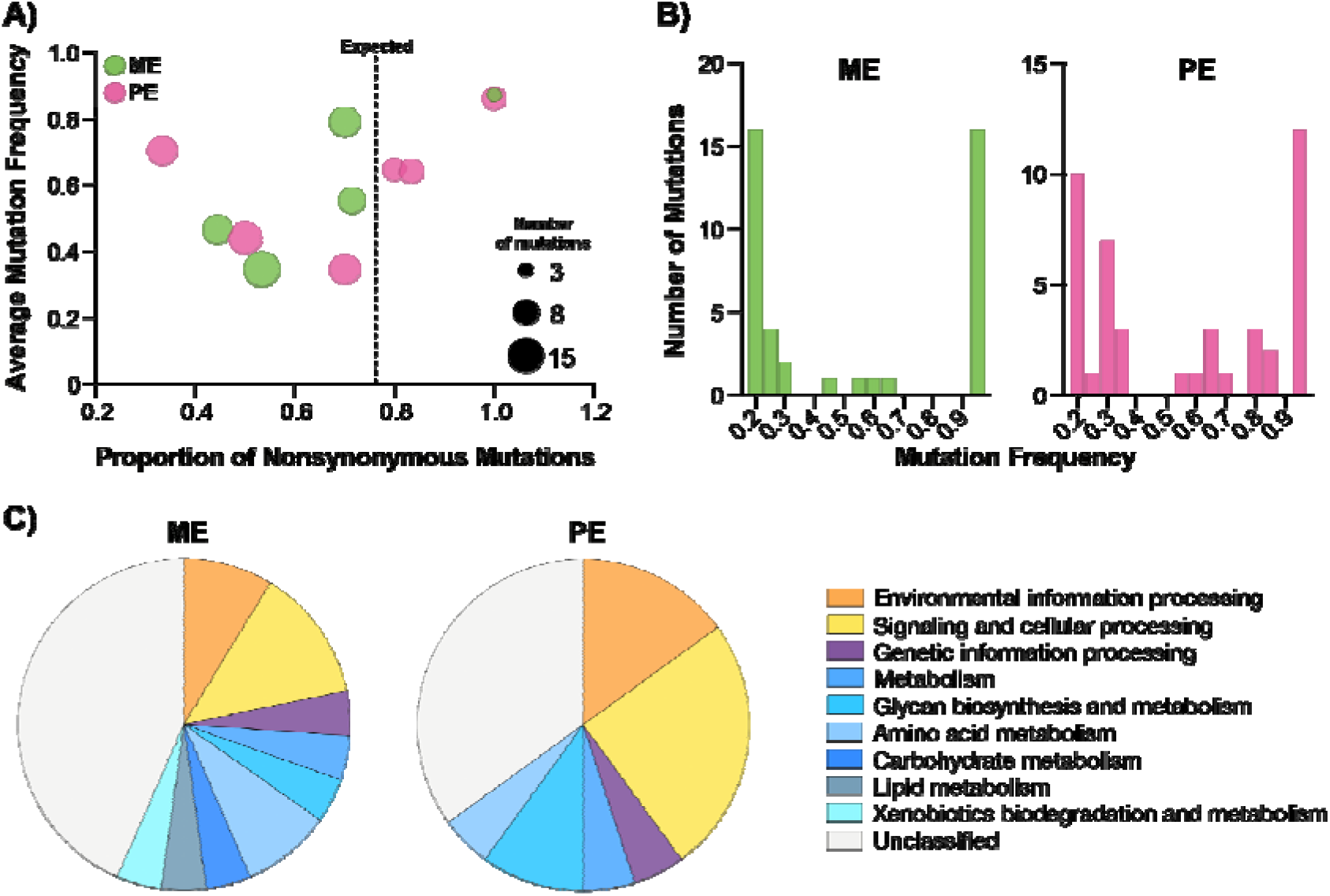
High-frequency (≥0.20) mutations present in the evolved populations. **A)** Mutation frequency and proportion of nonsynonymous mutations across all evolved populations. Scatter plots show the average mutation frequency (y-axis) versus the proportion of nonsynonymous and nonsense mutations (x-axis) for each evolved population at D60. Circle size represents the total number of mutations detected per population. The vertical dashed line represents the number of nonsynonymous + nonsense mutations expected by random chance. **B)** Number of mutations detected at different frequencies between 0.2 and 1 in ME and PE populations. Each bar represents the lower bound of each 0.05 interval (e.g., 0.2 = 0.2–0.24; 0.95 = 0.95–1.0). **C)** Functional annotation of coding-region mutations assigned to KEGG Orthology (KEGG) functions in media-evolved (ME) and predator-evolved (PE) lines; noncoding mutations are not shown. See Dataset S2 for a complete list of identified mutations.

**Figure 3.**
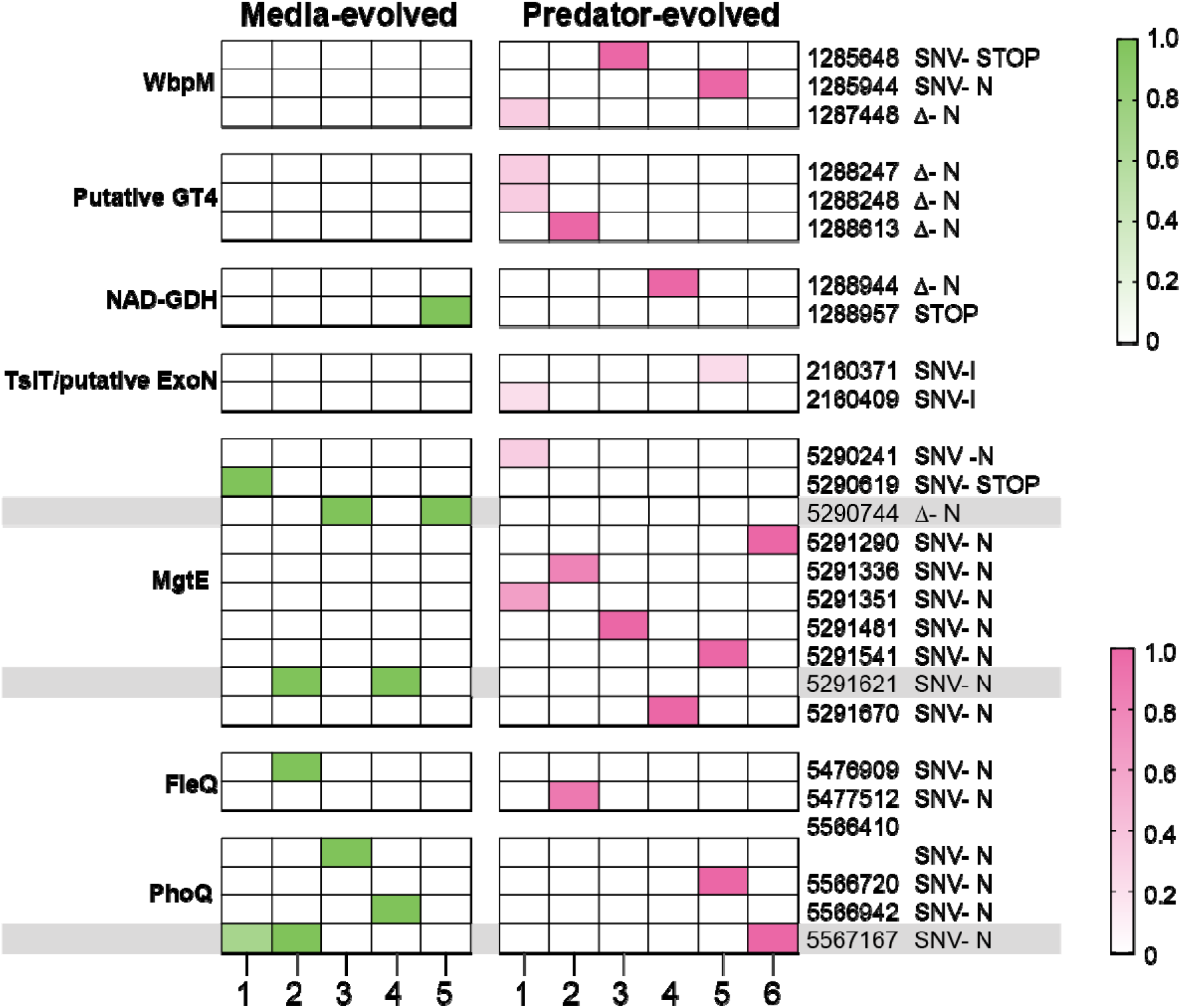
Parallel mutations detected in the evolved populations. Parallel mutations (same gene or same site) present at a frequency of at least 20% within one population. Bars on the right show what condition the mutation is present in (green = ME, pink = PE, gray = both). Full details of mutation frequencies and mutation types are provided in Dataset S2.

### High-frequency mutations reveal adaptive evolutionary dynamics

Restricting analyses to mutations present at ≥0.20 frequency in at least one population yielded 81 mutations, with between 3 and 15 mutations per population (Figure 2A; Dataset S2). On average, populations contained 8.3 mutations, and mean mutation frequencies ranged from 0.35 to 0.87.

The proportion of nonsynonymous mutations varied across populations (Figure 2A). Most ME populations and approximately half of the PE populations exhibited fewer nonsynonymous mutations than expected under a random model, consistent with purifying selection acting to remove deleterious variants. By contrast, the remaining PE populations exhibited a higher proportion of nonsynonymous mutations than expected, suggesting positive selection promoting the spread of beneficial mutations in these populations (Figure 2A).

The distribution of filtered mutation frequencies, calculated using 0.05 frequency bins, was bimodal, with prominent peaks at 0.20–0.24 and ≥0.95 (Figure S2). Mutations present at frequencies ≥0.95 were classified as fixed. Differences emerged between treatments. PE populations exhibited a broader distribution of mutation frequencies, with a substantial proportion of mutations at intermediate frequencies, whereas ME populations displayed a pronounced bimodal distribution, with mutations clustering around ~0.2–0.3 and ≥0.95 (Figure 2B). These patterns suggest distinct evolutionary dynamics: PE populations likely maintain a larger number of competing lineages, consistent with clonal interference, whereas ME populations show evidence of selective sweeps, as indicated by the enrichment of high-frequency mutations, while also retaining a subset of lower-frequency variants, indicating that competition among mutations is not absent.

To further characterize the functional basis of adaptation, we assigned KEGG orthology annotations to genes harboring mutations. Functional categorization revealed differences between treatments (Figure 2C). In both ME and PE populations, mutations were distributed across multiple functional classes, with a substantial proportion remaining unclassified. However, PE populations showed a relative enrichment of mutations in categories associated with environmental information processing and signaling and cellular processes, whereas ME populations exhibited a broader distribution across metabolic pathways. These differences suggest that adaptation in PE populations may preferentially target pathways involved in environmental sensing and interaction, while adaptation in ME populations is more broadly distributed across metabolic functions.

We observed strong signatures of parallel evolution, with multiple independent mutations targeting a limited set of genes across replicate populations (Figure 3). Notably, the magnesium transporter *mgtE* and the sensor kinase *phoQ* were recurrently mutated across both treatments, frequently reaching high frequency or fixation. However, the extent and pattern of parallelism differed between treatments.

To quantify parallel evolution, we tested whether mutations occurred in the same genes or at identical genomic positions more often than expected under a null model. Expected values were generated using simulations that accounted for the number of mutations, nucleotide-specific mutability, and mutation spectra observed in each population (see Methods).

We observed a substantial excess of gene-level parallelism across all treatments (Figure 3, Dataset S2). In ME populations, 12 mutations occurred in two genes (*phoQ* and *mgtE*), compared to an expected 0.1361 under the null model. Similarly, PE populations exhibited 26 mutations across four genes and one intergenic region, compared to an expected 0.12. Across both treatments combined, including genes shared between ME and PE, 44 mutations occurred in recurrently targeted loci, compared to an expected 0.326, representing an approximately 80–200-fold enrichment over random expectation.

At the site level, parallelism was less pronounced than at the gene level but remained elevated above random expectation. In ME populations, six identical-site mutations were observed (expected 0.011), whereas no identical-site mutations were observed within PE populations (expected 0.021). In addition, three identical-site mutations were shared between ME and PE populations (Figure 3, Dataset S2), compared to an expected 0.034.

Together, these results demonstrate that adaptation in this system is both genetically constrained and shaped by population-level processes. Selection repeatedly targets a limited set of genes, and even specific nucleotide positions, indicating highly constrained adaptive solutions. At the same time, evolutionary dynamics differ between treatments: ME populations exhibit more efficient fixation of beneficial mutations consistent with selective sweeps, whereas PE populations maintain greater genetic diversity likely due to clonal interference among competing lineages.

### Genomic and phenotypic characterization of evolved isolates

A single isolate was collected from each population (ME, n = 5; PE, n = 6). Analysis of each isolate revealed between three to seven mutations per genome, and the majority of mutations fixed in each population were present in the corresponding isolate. In total, 42 mutations were identified across 22 genes and one intergenic region (Figure 4A, Dataset S3). Mutations in the isolates were strongly biased toward nonsynonymous and nonsense changes, suggesting that many were adaptive (Figure 4B).

**Figure 4.**
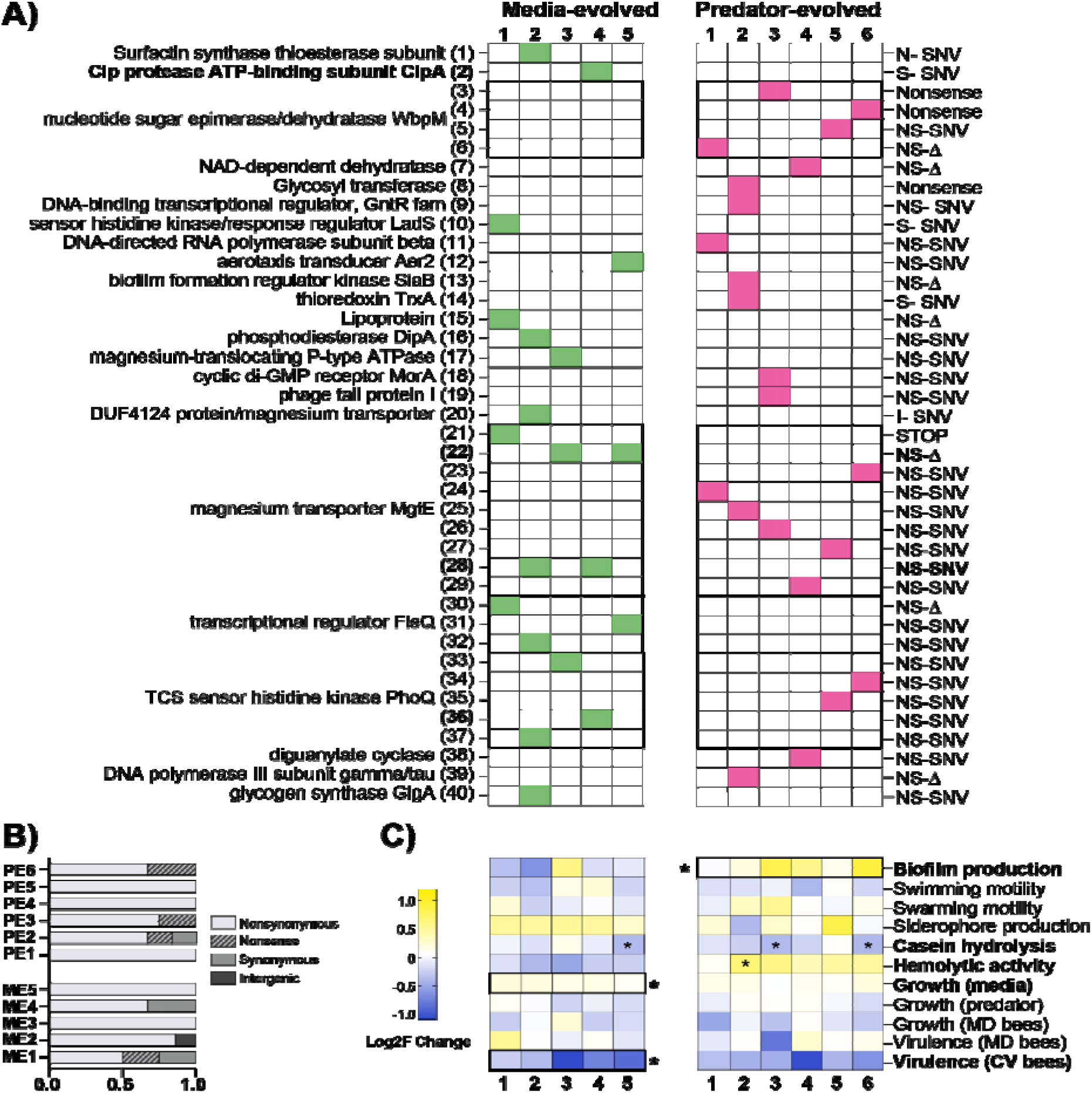
Genotypic and phenotypic changes in evolved isolates. **A)** Mutations identified in evolved isolates, highlighting site-specific parallel mutations (bold font) and gene-specific parallel mutations (outlined boxes). NS, nonsynonymous; S, synonymous; I, intergenic; STOP, nonsense; SNV, single-nucleotide variant; Δ, deletion; +, insertion. For a list of all mutations identified in the evolved isolates, see Dataset S3. **B)** Proportions of mutation types (NS, S, I, STOP) present in each isolate. **C)** Log_2_ fold change calculated from the mean of all replicates and trials for each phenotype measured in evolved isolates relative to the ancestral strain. Yellow indicates an increase, blue indicates a decrease, and white indicates no change. Asterisks within boxes indicate significant differences when individual isolates were compared to the ancestor, whereas asterisks in the margin indicate significant differences when the mean of all ME lines or all PE lines was compared to the ancestral mean. Bold text denotes phenotypes showing significant changes relative to the ancestor. Statistical significance was determined using a Kruskal–Wallis test with Dunn’s multiple comparison test (P < 0.05). Multiple-comparison corrections were applied independently within each phenotypic assay, rather than across the full set of measured traits. Full data for all replicates, including non-log_2_ fold-change values, are shown in Figures S4 and S5.

Reflecting patterns observed at the population level, several instances of parallel evolution were detected across isolates. Among the ME isolates, two distinct site-specific parallel mutations in *mgtE* were identified in ME2 and ME4 and in ME3 and ME5. Additionally, a gene-specific mutation in *fleQ*, unique to the ME lines, was observed in ME1, ME2, and ME5 (Figure 4A, Dataset S3). In the PE isolates, a gene-specific mutation in *wbpM*, unique to the PE lines, was identified in PE1, PE2, PE5, and PE6. Finally, gene-specific mutations in *mgtE* and *phoQ* were shared between ME and PE isolates (Figure 4A, Dataset S3).

To assess the impact of predator-driven evolution on *P. aeruginosa*, we performed phenotypic assays on a single isolate from each evolved population. Virulence-associated traits, including biofilm production, motility, siderophore production, casein hydrolysis, and hemolytic activity, were measured (Figure 4C, Figure S3 A–F). Fitness was evaluated in two different conditions: growth after 24 h in media and in coculture with *T. thermophila* (Figure 4C, Figure S3 G–H). All assays were performed in at least triplicate. Because the data did not consistently meet assumptions of normality, we used nonparametric statistical tests, which are more robust to violations of distributional assumptions but may have reduced statistical power, potentially limiting the detection of modest but biologically relevant differences. Statistical comparisons were performed both at the level of individual isolates and at the group level, where all isolates from each treatment (ME or PE) were pooled and compared to the mean of the ancestral strain. Statistical corrections for multiple comparisons were applied independently within each assay type rather than across the full set of assays.

Phenotypic responses were heterogeneous across isolates, with few consistent directional changes. Specifically, isolates PE3, PE6, and ME5 exhibited reduced casein hydrolysis, indicating decreased extracellular protease activity involved in protein degradation (*P* = 0.0078, 0.0059, and 0.0076, respectively; Figure 4C, Figure S3E), whereas isolate PE2 showed significantly increased hemolytic activity compared to the ancestral strain (*P* = 0.0362, Figure 4C, Figure S3F). When comparing the mean of all PE or ME isolates to the ancestor, PE isolates showed higher biofilm production (*P* = 0.0362; Figure 4C, Figure S3A) and ME isolates exhibited increased growth in media (*P* = 0.0054; Figure 4C Figure S3G).

Although not statistically significant, additional patterns were observed. Swimming activity (Figure 4C, Figure S3B) decreased in most D60-evolved lines, with the exception of ME3 and ME4. Swarming activity (Figure 4C, Figure S3C) was generally reduced in ME isolates, except ME1, but increased in most PE isolates relative to the ancestor. For biochemical traits, most evolved isolates produced slightly more siderophore (Figure 4C, Figure S3D), with PE2 as the exception. Protease activity decreased on average in both ME and PE lines. Casein hydrolysis decreased in all ME lines, whereas PE lines exhibited an increase (Figure 4C, Figure S3E).

To evaluate how predation impacted *P. aeruginosa* virulence, we used a tractable invertebrate host model, the honey bee (*Apis mellifera*). This system was selected because the ancestral strain exhibited moderate virulence in bees, allowing us to assess whether evolved isolates showed increased virulence in a model that supports large numbers of replicate hosts. We performed two types of assays: one in conventional (CV) honey bees, which were healthy adults with an intact microbiome, and another in microbiota-depleted (MD) bees. The MD bees were newly emerged (~1 day old) adults of the same age.

All ME and PE isolates showed a decrease in virulence on average in CV bees compared to the ancestor, although these differences were not significant after correction for multiple comparisons. When analyzed at the group level, ME isolates exhibited a significant reduction in virulence, whereas PE isolates did not differ significantly from the ancestor (Figure 4C, Figure S4A).

In MD bees, virulence responses were more variable, with some isolates showing slight increases and others slight decreases relative to the ancestor. ME1, ME4, PE4, and PE6 exhibited slight increases in virulence, while all other isolates showed slight decreases; however, none of these changes were statistically significant. When analyzed by treatment, neither ME nor PE isolates differed significantly in virulence compared to the ancestor (Figure 4C, Figure S4B).

To determine whether the attenuated virulence was due to reduced proliferation in the honey bee gut, we performed growth assays for each isolate in MD bees (Figure 4C, Figure S4C). While the mean growth levels showed trends that loosely resembled the virulence patterns observed in MD bees (Figure 4C), no statistically significant differences in growth were detected for either ME or PE isolates. Instead, both groups exhibited substantial isolate-to-isolate variation. In each group, three of the six replicates showed a slight increase in growth (ME2, ME3; PE2, PE4, PE5), whereas the remaining three showed a slight decrease (ME1, ME4, ME5; PE1, PE3, PE6) relative to the ancestor.

Each isolate and the ancestor were grown individually and standardized to 1 OD. Equal volumes of each were then combined to generate a 0.11 mL mixed inoculum, which was subsequently cultured for 24 h in media, in the presence of *T. thermophila*, or in honey bees (Figure 5). WGS metagenomic sequencing was then used to determine which strains outcompeted the others in each condition. In the presence of the predator, ME3 and ME4 outcompeted all other lines. In media, ME2, ME3, and ME4 outcompeted the other isolates and the ancestor. In honey bee competitions, ME4 prevailed over all other isolates and the ancestral strain (Figure 5).

**Figure 5.**
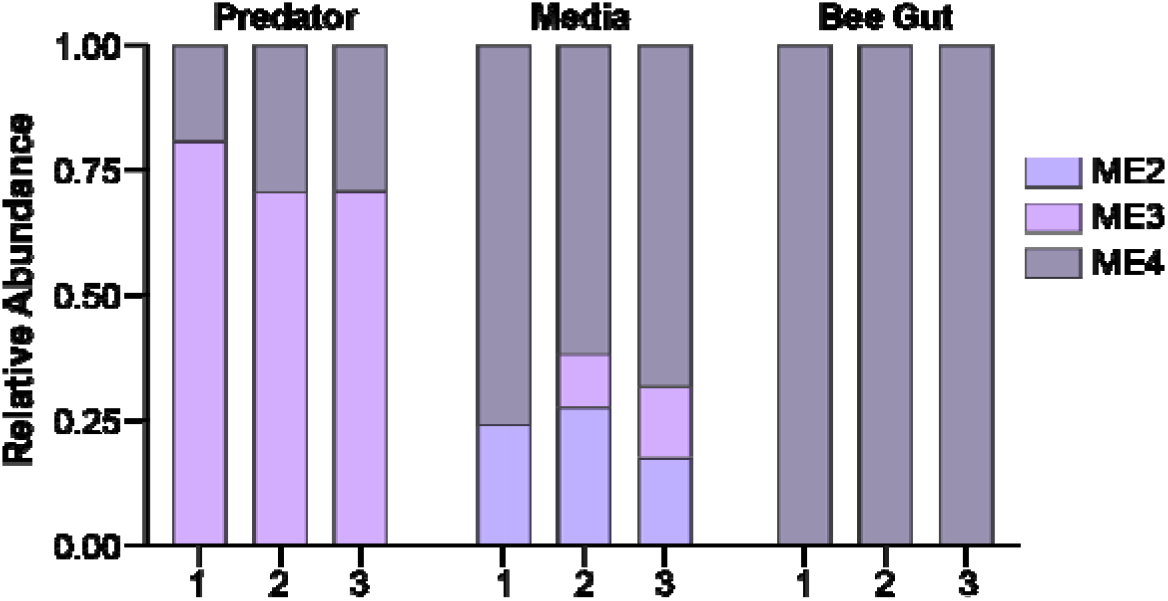
**Competition assays with the evolved isolates and the ancestral strain** in the presence of a predator, in media, or in honey bee host. Each assay was performed in triplicate.

The observed competition outcomes are highly deterministic as the probability that the assay would randomly result in the same combinations of strains three times independently is extremely low (*p*~10-8 assuming that up to three strains can colonize the same environment; See Methods Section).

## Discussion

Predator-mediated selection is widely recognized as a major evolutionary force shaping bacterial phenotypes, genome content, and ecological interactions (Buckling and Rainey 2002; Matz and Kjelleberg 2005). In this study, we examined how exposure to the ciliate predator *T. thermophila* influences the evolutionary dynamics, genetic architecture, and virulence of *P. aeruginosa*. By combining experimental evolution with whole-genome sequencing and phenotypic assays, we show that while adaptation is strongly constrained at the genetic level, predation fundamentally alters the dynamics of evolution, promoting genetic diversity and modifying how environmental adaptation translates into host-associated outcomes.

A central finding of this work is that predation reshapes the mode of adaptation rather than simply redirecting its genetic targets. In the absence of predators, ME populations were characterized by a higher proportion of near-fixed mutations and a pronounced bimodal distribution of allele frequencies, consistent with repeated selective sweeps in a relatively stable environment. By contrast, PE populations exhibited a broader distribution of allele frequencies, with a higher prevalence of intermediate-frequency mutations, indicative of clonal interference and the coexistence of competing lineages. These patterns are consistent with theoretical expectations that more complex and heterogeneous environments, particularly those involving biotic interactions, expand the adaptive landscape and maintain multiple competing solutions (Rainey and Travisano 1998; Desai et al. 2007; Amicone and Gordo 2021). In this context, predation acts less as a directional selective pressure and more as a source of ecological complexity.

Despite these differences in evolutionary dynamics, adaptation remained highly constrained at the genetic level. We observed strong enrichment of parallel evolution, with repeated targeting of genes such as *mgtE* (a dual-function magnesium transporter linked to type III secretion system regulation; Anderson et al. 2010; Coffey et al. 2014) and *phoQ* (part of a two-component regulatory system; Gellatly et al. 2012) across replicate populations. This pattern indicates that selection consistently favors a limited set of functional pathways, even under distinct environmental conditions. However, mutations within these shared targets frequently occurred at different genomic positions, suggesting that while evolution is predictable at the level of gene targets, it remains flexible at the level of specific molecular changes. Such divergence within shared loci may generate distinct phenotypic outcomes, reinforcing the idea that adaptation reflects a balance between constraint and contingency.

Differences in the functional distribution of mutations between treatments further suggest that predator-mediated selection alters the targets of adaptation. While mutations in both ME and PE populations spanned multiple functional categories, PE populations showed a relative enrichment of mutations in pathways associated with environmental information processing and signaling and cellular processes. By contrast, mutations in ME populations were more broadly distributed across metabolic functions. These patterns are consistent with previous work showing that bacterial responses to predation involve traits related to environmental sensing, surface modification, and interaction with biotic factors (Hahn et al. 2000; Matz et al. 2004; Matz et al. 2008; Jousset et al. 2009). At the same time, the substantial overlap in mutations between ME and PE populations indicates that adaptation to the abiotic growth environment remained a dominant selective force. Together, these results suggest that predator-driven selection shifts the functional targets of adaptation without fully redirecting them, promoting divergence in evolutionary trajectories while maintaining a shared core of adaptive responses.

These dynamics are also shaped by the balance between selection and drift under the experimental design. Daily transfers imposed repeated bottlenecks, which likely reduced effective population size and increased the role of stochastic processes (Wahl and Gerrish 2001; Good et al. 2012). Such effects may have been especially pronounced in PE populations if predator exposure reduced bacterial abundance or increased temporal variability in growth. Under these conditions, both clonal interference and drift may contribute to the persistence of intermediate-frequency mutations and the reduced prevalence of fixation. Nevertheless, the strong excess of parallel evolution at both the gene and site levels indicates that selection remained a dominant force, acting within a framework where demographic and ecological factors modulate evolutionary outcomes. We note that detection of low-frequency variants in metagenomic datasets can be influenced by coverage variation, read mapping ambiguity, and localized sequencing error hotspots, which we sought to mitigate through stringent filtering and exclusion of variants also detected in the ancestral population. These considerations primarily affect very low-frequency calls, whereas the overall patterns of parallel evolution remain robust and consistently supported across populations.

Although genomic signatures of adaptation were pronounced, phenotypic changes were generally modest and often not statistically significant. This apparent disconnect likely reflects the polygenic and pleiotropic nature of many bacterial traits (Desai et al. 2007; Kryazhimskiy et al. 2014; Good et al. 2017) as well as the relatively small effect sizes of individual mutations. Because the functional consequences of specific mutations are often difficult to predict, particularly in regulatory or multifunctional genes, linking individual genetic changes to phenotypic outcomes remains challenging. Instead, adaptation in this system likely reflects the cumulative effects of multiple mutations acting across interconnected pathways. Consistent with this, mutations in loci such as *mgtE, wbpM, PhoQ*, and other regulatory genes are known to influence a range of traits, including motility, growth, and virulence-related processes (Hickman and Harwood 2008; Anderson et al. 2009; Gellatly et al. 2012; Davis et al. 2013; Ravichandran et al. 2015), suggesting that adaptation proceeds along constrained phenotypic trajectories shaped by pleiotropy and epistasis.

Within this framework, our virulence results highlight the importance of evolutionary dynamics in shaping host-associated outcomes. The coincidental evolution hypothesis predicts that traits selected during interactions with environmental predators may enhance virulence in animal hosts (Levin 1996; Matz and Kjelleberg 2005; Adiba et al. 2010; Friman and Buckling 2014). In our system, however, virulence outcomes were variable rather than directional. Most evolved isolates exhibited modest reductions in virulence, while a subset showed slight increases depending on host context. These findings suggest that predator-mediated selection does not uniformly drive virulence in a single direction, but instead generates a spectrum of phenotypic outcomes.

One explanation for this variability is that the increased genetic diversity and clonal interference observed in PE populations maintain multiple competing adaptive strategies, some of which may enhance virulence while others reduce it. In this view, predator-driven environments may preserve phenotypic variation rather than pushing populations toward a single virulence optimum. By contrast, adaptation in more stable environments, such as the ME treatment, may promote more consistent directional changes through repeated selective sweeps, potentially leading to more uniform outcomes such as reduced virulence. More broadly, these results suggest that the relationship between environmental selection and virulence depends not only on the traits under selection but also on the underlying evolutionary dynamics.

The use of the honey bee model is best understood within this conceptual context. Our goal was not to replicate the selective pressures imposed by *T. thermophila* but to test whether adaptation to a eukaryotic predator alters pathogenic potential in a distinct eukaryotic host. This approach directly addresses the logic of coincidental virulence by asking whether traits favored in one ecological interaction influence outcomes in another. Honey bees provide a tractable and scalable *in vivo* system in which the ancestral strain exhibits moderate virulence, allowing detection of both increases and decreases in pathogenicity across many replicates. At the same time, because *P. aeruginosa* is not a natural pathogen of bees, the observed virulence patterns should be interpreted as relative changes in pathogenic potential rather than host-specific adaptation.

The substantial overlap in adaptive mutations between ME and PE populations also highlights an important consideration for experimental design. In the present study, ME lines adapted to the abiotic growth medium alone, whereas PE lines adapted simultaneously to both the medium and the predator. This difference in selective regimes may have contributed to the observed overlap in mutations as well as to potential trade-offs and clonal interference in PE populations, where competing beneficial mutations associated with abiotic and biotic adaptation could arise concurrently. Early adaptation to the abiotic growth medium likely contributed significantly to the observed evolutionary trajectories, potentially obscuring more subtle predator-specific effects. Pre-conditioning in the Neff medium prior to predator exposure could have helped reduce the contribution of abiotic adaptation and allowed clearer resolution of predator-driven responses. Additionally, the 60-day evolution period (~276 generations) may not have been sufficient for all adaptive mutations, particularly those with smaller fitness effects or those specific to predator interactions, to reach high frequency or fixation. Extending the duration of the experiment or allowing populations to first equilibrate to the abiotic environment, could result in stronger genomic signals and more pronounced phenotypic divergence. However, because abiotic and biotic selection pressures are intertwined in natural environments, the combined effects observed here may also reflect the ecological reality faced by opportunistic pathogens.

Competition assays further illustrate the context dependence of adaptive outcomes. ME isolates consistently outcompeted both the ancestral strain and PE isolates across multiple environments, including in the presence of predators. While this may appear counterintuitive, it suggests that traits conferring resistance to predation do not necessarily translate into higher competitive fitness in mixed populations. Predator-driven adaptation may involve trade-offs, such as reduced growth rate or altered resource allocation, that diminish performance when competing directly with faster-growing strains. These results reinforce the idea that different components of fitness, such as predator resistance, growth, and competitive ability, can evolve semi-independently and yield environment-specific advantages.

In summary, our findings demonstrate that adaptation in *P. aeruginosa* is shaped by a combination of genetic constraint and environment-dependent evolutionary dynamics. Predator-mediated selection increases ecological complexity, promotes genetic diversity, and alters the balance between selection, drift, and clonal interference, while leaving core adaptive targets largely unchanged. Rather than driving a consistent increase in virulence, predation appears to broaden the range of possible host-associated outcomes by maintaining multiple evolutionary trajectories. These results refine our understanding of how ecological interactions shape pathogenic potential and highlight the importance of considering evolutionary dynamics, in addition to selective pressures, when predicting the emergence of virulence.

## Methods

### Experimental evolution

The ancestral *P. aeruginosa* “PAO1” strain we used for our experiments was purchased from Carolina Biological Supply Company and evolved either in media alone (ME) or in the presence of the protist predator *T. thermophila* strain SB210 (PE). *T. thermophila* was obtained from the Tetrahymena Stock Center at Cornell University. Each evolution treatment included six replicate lines.

Prior to starting the evolution experiment, *T. thermophila* and ancestral *P. aeruginosa* were cultured individually for 72 h and 48 h, respectively, at 35°C in Neff media. To initiate each evolved line, 1 OD of *P. aeruginosa* was resuspended in fresh Neff media, and 0.12 mL was added to either 3 mL of Neff media alone (ME lines) or 3 mL of Neff media containing *T. thermophila* (PE lines). The starting ratio of bacteria to predator was determined in preliminary trials to allow both organisms to persist in coculture for 24 h.

Experimental cultures (3 mL) were incubated at 35°C with shaking. Every 24 h for 60 days (~276 generations), cultures were vortexed and 4% of each line was transferred into fresh Neff media (ME lines) or new axenic *T. thermophila* cultures (PE lines). Prior to each transfer, *T. thermophila* viability and density were assessed using light microscopy, and *P. aeruginosa* growth was confirmed by plating on LB agar. Throughout the experiment, both organisms were consistently present, and no noticeable changes in population size were observed based on daily growth and plating. On day 60, agar plates were inoculated from each culture, and one colony was randomly selected from each ME (ME1–6) and PE population (PE1–6). Each colony was grown in 3 mL LB broth for 48 h at 35°C to obtain pure cultures. A 200 μL aliquot of each culture was frozen at −80°C in 20% glycerol. These 12 isolates were subsequently sequenced and used for phenotypic assays, including virulence in bees, growth assays, biochemical tests, biofilm production, and competition experiments.

### DNA extraction and sequencing

Cultures of the evolved populations (ME1–6 and PE1–6), evolved isolates (ME1–6 and PE1–6), and the ancestral strain were diluted to 1 OD, and DNA was extracted using the Zymo Quick-DNA™ Miniprep Plus Kit (D4069). Libraries for short-read sequencing were prepared using the Illumina Nextera DNA Flex Library Prep Kit (20018704). The ancestral genome was sequenced using both long-read (Oxford Nanopore MinION) and short-read (Illumina iSeq100) technologies. Evolved populations and isolates were sequenced using short-read Illumina platforms (iSeq100 or MiSeq100) with 2 × 150 bp paired-end reads.

### Genome assembly and analysis

The Illumina and Nanopore reads of the ancestral genome were combined to generate a hybrid assembly using hybridSPAdes v3.15.3 (Antipov et al. 2016). Assembled scaffolds were annotated with Bakta (Schwengers et al. 2021) on the Department of Energy Systems Biology Knowledgebase (KBase) platform. Genome sequence comparisons were performed between the ancestral strain and the three closest related strains, identified based on BLAST searches against the NCBI database. Average nucleotide identity values were calculated using FastANI v1.32 (Jain et al. 2018). A pangenome analysis was conducted using Panaroo v1.5.2 (Tonkin-Hill et al. 2020) with the following parameters: an initial clustering threshold of 0.98 using CD-HIT, a family threshold of 0.7 for secondary clustering of orthologous genes into gene families, a length difference percentage cutoff of 0.80, a core gene threshold of 1 (genes present in all genomes), strict clean mode, and removal of invalid genes. All short reads of the ancestral strain were then mapped back to the consensus genome using breseq v0.39.0 (Deatherage and Barrick 2014). Any variants identified during this mapping were considered sequencing or assembly artifacts and were excluded from the set of candidate mutations in the evolved populations and isolates.

Sequencing reads from evolved populations and isolates were trimmed using Trimmomatic v0.40 (Bolger et al. 2014) and mapped to the assembled ancestral genome using breseq v0.39.0 (Deatherage and Barrick 2014). Mutations were called using a base-quality cutoff of Q≥30, requiring at least two mutation-supporting reads per strand in polymorphism mode, and a minimum frequency threshold of 0.05. Identified mutations were compared to the ancestral strain using gdtools (Deatherage and Barrick 2014). Additional filtering removed mutations present at frequencies ≤0.2. However, if a parallel mutation at the same genomic position reached a frequency ≥0.2 in any population, all mutations at that position were retained. The functional characterization of genes was predicted using KOALA KEGG Orthology and Links Annotation (Kanehisa 2000; Kanehisa et al. 2016).

#### Clonal deconvolution

Potential contamination between evolved lines was assessed by constructing pseudoclones. For each population, allele variants were grouped into frequency bins (<10%, 10–20%, 20–30%, …, >90%), and variants within each bin were merged to generate a corresponding pseudoclone genome by introducing the observed mutations in each bin into the reference genome. The resulting pseudoclone genomes were aligned using MUSCLE v5 (Edgar 2022). Phylogenetic analysis of the aligned pseudoclones was performed in SeaView v5.1 (Gouy et al. 2009) using PhyML v3.0 (Guindon et al. 2010) with the GTR model. Trees were visualized using FigTree v1.4.4. (Rambaut 2018).

### Phenotypic assays

#### Growth in media

Cultures of evolved populations (ME1–6 and PE1–6), evolved isolates (ME1–6 and PE1–6), and the ancestral strain were diluted to 1 OD. For each isolate, 0.12 mL of culture was added to 3 mL fresh Neff media in triplicate and incubated at 35°C with shaking at 110 rpm. After 24 h, cultures were diluted and plated on cetrimide agar in duplicate. Plates were incubated for 48 h at 35°C, and colonies were counted to calculate CFUs.

#### Growth in the presence of *T. thermophila*

Axenic 48-h cultures of *T. thermophila* in Neff media were inoculated with 120 μL of 1 OD cultures of each isolate in triplicate. Cultures were incubated for 20 h at 35°C and 110 rpm, diluted, and plated on cetrimide agar in duplicate. Plates were incubated for 48 h at 35°C before counting colonies and calculating CFUs.

#### Growth in-host

Pupae were collected 2–3 days before emergence and kept in cages with sterile sugar syrup (SSS) and sterile pollen. Newly emerged adult bees were hand-fed 10 μL of either SSS, the ancestor in SSS, or an evolved isolate in SSS (5 bees per cup, 15 bees per treatment). Bees were maintained at 35°C and 95% humidity. After 20 h, guts were collected, stored in 20% glycerol at −80°C, and dilutions up to 10^−6^ were performed and plated on cetrimide agar in triplicate. Plates were incubated 48 h at 35°C, colonies were counted, and CFUs/mL were calculated based on the dilution factor.

#### Biofilm assay

Biofilm formation was assessed in triplicate. First, 5 μL of 1 OD cultures were added to 150 μL Neff media in 96-well microplates with an additional sterile nonskirted plate immersed into the wells to provide a bigger surface attachment; this setup was incubated at 35°C for 24 h without shaking. Wells and the immersed plate were washed twice with sterile ddH_2_O, stained with 150 μL of 0.1% crystal violet for 15 min, washed again, and air-dried. Biofilms in both microplates (initial and immersed) were separately solubilized with 150 μL of 30% acetic acid, and absorbance was measured at 490 nm. The measurements were combined to obtain a final reading. Three independent assays were performed, each with four replicates.

#### Casein hydrolysis

Milk agar plates were prepared by mixing 2.5 g yeast extract and 7.5 g agar in 350 mL water and 15 g milk powder in 150 mL water; both solutions were autoclaved, cooled, mixed, and poured into plates. Sterile discs were inoculated with 4 μL of 1 OD bacterial culture and incubated at 37°C for 48 h. To assess protease activity, the plates were photographed and analyzed using ImageJ by calculating the diameter of the activity and subtracting the 4-mm diameter of the disc.

#### Siderophore production

Chromeazurol S (CAS)–Cetyltrimethylammonium bromide (CTAB) agar plates were prepared by dissolving 0.065 g CAS in 50 mL water and 0.0729 g CTAB in 40 mL water. Those two solutions were then mixed with 10 mL of 1 mM FeCl_3_ hexahydrate and 10 mM HCl solution and added to LB agar medium, followed by pH adjustment to 6.8 and autoclaving. Sterile discs were placed and inoculated with 4 μL of 1 OD bacterial culture and incubated at 37°C for 48 h. For analyzing siderophore production, plate photographs were uploaded to ImageJ to measure the diameter of the siderophore production minus the diameter of the disc (4 mm).

#### Hemolytic activity

Blood agar plates (LB + 5% sheep blood) were inoculated with 4 μL of 1 OD bacteria and incubated at 37°C for 48 h before assessing hemolysis, as previously described in the siderophore production protocol.

#### Swimming and swarming motility

Swim plates contained peptone (5 g), yeast extract (3 g), and 0.3% agar per liter of water; swarm plates used 0.5% agar. Plates were autoclaved, poured, and allowed to solidify for up to 24 h. The plates were stab-inoculated with 2 μL of 1 OD culture and incubated at 37°C for 48 h before measuring motility by measuring the diameter of the motion in photographs uploaded to ImageJ.

### Virulence assay in honey bees

#### Conventional bees

Pure 1 OD cultures of each evolved isolate and the ancestor were centrifuged (10,000 rpm × 2 min) and individually resuspended 1:1 in SSS. Adult worker bees (*Apis mellifera*) were collected from a single hive at the Phytotron, NCSU, Raleigh, NC, and immobilized at 4°C. Bees were randomly distributed into groups and exposed via the immersion method (~10 μL per bee) to one of three treatments: 1) sterile SSS only, 2) the ancestor in SSS, or 3) each evolved isolate in SSS. Bees were maintained in cup cages (20 bees/cup; 100 bees/treatment) under hive-like conditions (35°C, 95% humidity). Mortality was recorded daily for 5 days. Two independent assays were performed, each with five replicates, totaling 200 bees per isolate.

#### Microbiota-depleted bees

Each evolved isolate was first cultured to 1 OD_600_, followed by centrifugation to pellet the bacteria and resuspension of the pellet in 1:1 SSS. Newly emerged honey bees (*A. millifera*) were immobilized at 4°C and randomly distributed in different 50-mL Falcon tubes. Ten bees were added to a single tube, in which isolate-containing SSS was added (10 μL per bee) for immersion treatment. Every isolate, ancestor, and control treatment were performed in triplicate (i.e., 10 bees × 3 = 30 bees per treatment). The treated bees were then maintained in cup cages under hive-like conditions (35°C and 95% humidity) for 5 days, and mortality was recorded. Two independent assays were performed, totaling 60 bees per isolate.

### Competition assays

#### In-host competition

Pupae were collected 2–3 days prior to emergence from a single hive at the Phytotron (NCSU) and maintained in cages with SSS and sterile pollen. After emergence, cultures of each evolved isolate and the ancestor were normalized to OD_600_ = 1, centrifuged (10,000 rpm for 2 min), and resuspended in SSS. For inoculation, strains were pooled such that the ancestor and each evolved isolate (ME1–6 and PE1–6) were present in equal proportions. A total of 25 bees were hand-fed 10 µL of the bacterial suspension in SSS and maintained in cup cages (5 bees per cup; 5 cups total) at 35°C and 95% humidity. After 20 h, bee guts were collected under sterile conditions and stored in 20% glycerol at –80°C. Although all bees were sampled, three individual bee gut samples (each from a different cup cage) were selected for downstream analysis. These samples were plated on cetrimide agar in triplicate, incubated for 24 h, and ~300 colonies per sample were pooled in 1× PBS. DNA was extracted from the pooled colonies and analyzed by WGS metagenomic sequencing (2×150 PE reads on Illumina miSeq100). Reads were mapped to the ancestral genome using breseq v0.39.0 (Deatherage and Barrick 2014), and gdtools (part of the breseq pipeline) was used to compare each sample to mutations present in the ancestor and the individual evolved isolates (ME1–6 and PE1–6). The relative frequency of each strain was estimated as the average frequency of mutations shared between the isolate and the sample.

#### Competition in the presence of *T. thermophila*

Five replicate tubes containing 3 mL of Neff media were inoculated with 0.11 mL of a 1 OD mixture of ancestor and evolved isolates (prepared by combining 10 µL of each isolate standardized to 1 OD). Cultures were incubated for 20 h at 35°C with shaking at 110 rpm. Three replicate samples were diluted and plated in triplicate on cetrimide agar, incubated for 48 h at 35°C, and ~300 colonies per sample were collected in 1 mL of 1× PBS. DNA was extracted from the pooled colonies, sequenced, and analyzed following the protocol stated above for **in-host competition**.

#### Competition in media

Five replicate tubes containing 3 mL of Neff media were inoculated with 0.11 mL of a 1 OD mixture of ancestor and evolved isolates (prepared by combining 10 µL of each isolate standardized to 1 OD). After 20 h of incubation at 35°C and 110 rpm, three of the replicate cultures were centrifuged at 10,000 rpm for 2 min. Supernatants were discarded, and pellets were resuspended in 200 μL of 1× PBS. DNA was extracted from the pooled colonies (~300 colonies per sample), sequenced, and analyzed following the protocol described above for **in-host competition**.

### Statistical analyses

All figures and statistical analyses were performed in R, python, or GraphPad Prism v10.5.0. Figures were visually improved using Adobe Illustrator.

#### Mutations

The relationship between the total number of mutations per evolved population and (i) average genome coverage and (ii) average number of mutations was assessed using simple linear regression (see Figure 1A–B).

#### Estimation of expected mutation classes and parallel evolution

We conducted simulations to infer i) the proportion or mutations expected in each sequence type (i.e., nonsynonymous, nonsense, synonymous, RNA gene, or intergenic) and ii) the proportion of parallel mutations expected to impact the same gene and the same site in different lines. To do so, we characterized the number of mutations observed in each line as well as the observed mutability of each nucleotide (how frequently a mutation affects a C, A, G or T) and the observed mutation spectrum (e.g., how frequently a C mutates into A, G or T). For each line, we then simulated the observed mutation patterns using the ancestral genome 10,000 times. Sequence annotations of the ancestor were used to infer which type of sequence was mutated. Parallel mutations were estimated by comparing the number of mutations affecting the same gene or the same site across the different lines in each replicate.

#### Phenotypic analyses

For all phenotypic assays, significance between each evolved line and the ancestral strain was assessed using a Kruskal–Wallis test, followed by Dunn’s multiple comparisons test, incorporating both technical and biological replicates (see Figure 4C, Figure S3 and Figure S4). In addition, the mean phenotypic values of all ME lines combined, and all PE lines combined were each compared to the ancestral mean using a Kruskal–Wallis test with Dunn’s multiple comparisons test to assess differences between evolved groups relative to the ancestor. Multiple-comparison correction was applied independently within each phenotypic assay, rather than across the full set of measured assays (see Figure 4C, Figure S3, and Figure S4).

#### Competition assays

the probability of randomly obtaining the same combination of strains in three independent assays was computed as 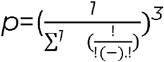, where *n* is the total number of strains and *k* the maximum number of strains that can co-colonize the same environment.

## Data availability

The complete genome of the ancestral strain has been deposited in NCBI under BioProject PRJNA1387885. All raw sequencing data for evolved populations and isolates have been deposited in the NCBI Sequence Read Archive (SRA) under BioProject PRJNA1390225. All other data are included in the main text and supplementary material. Bacterial strains and evolved populations are available from the authors upon reasonable request.

## Acknowledgements

We are grateful to Dr. Awa Diop for assistance in classifying the ancestral strain used in this study. We thank Dr. Parul Bardeskar and Dr. Michael Bland for assisting with honey bee experiments. We also thank Patrick Gallagher for careful reading of the manuscript and helpful editorial suggestions. This work was supported by the National Institutes of Health under grant 7R01GM145747 (to K.R.).

## References

Adiba, S., C. Nizak, M. van Baalen, É. Denamur, and F. Depaulis. 2010. From Grazing Resistance to Pathogenesis: The Coincidental Evolution of Virulence Factors. PLoS ONE 5. Public Library of Science. doi:10.1371/journal.pone.0011882.

Amaro, F., W. Wang, J. A. Gilbert, O. R. Anderson, and H. A. Shuman. 2015. Diverse protist grazers select for virulence-related traits in Legionella. The ISME Journal 9. Springer Nature: 1607. doi:10.1038/ismej.2014.248.

Amicone, M., and I. Gordo. 2021. Molecular signatures of resource competition: Clonal interference favors ecological diversification and can lead to incipient speciation*. Evolution 75. Oxford University Press: 2641. doi:10.1111/evo.14315.

Anderson, G. G., T. L. Yahr, R. R. Lovewell, and G. A. O’Toole. 2009. The Pseudomonas aeruginosa Magnesium Transporter MgtE Inhibits Transcription of the Type III Secretion System. Infection and Immunity 78. American Society for Microbiology: 1239. doi:10.1128/iai.00865-09.

Antipov, D., A. Korobeynikov, J. S. McLean, and P. A. Pevzner. 2015. hybridSPAdes: an algorithm for hybrid assembly of short and long reads. Bioinformatics 32. Oxford University Press: 1009. doi:10.1093/bioinformatics/btv688.

Barbosa, C., V. Trebosc, C. Kemmer, P. Rosenstiel, R. Beardmore, H. Schulenburg, and G. Jansen. 2017. Alternative Evolutionary Paths to Bacterial Antibiotic Resistance Cause Distinct Collateral Effects. Molecular Biology and Evolution 34. Oxford University Press: 2229. doi:10.1093/molbev/msx158.

Bolger, A., M. Lohse, and B. Usadel. 2014. Trimmomatic: a flexible trimmer for Illumina sequence data. Bioinformatics 30. Oxford University Press: 2114. doi:10.1093/bioinformatics/btu170.

Brown, S. P., L. L. Chat, and F. Taddéi. 2007. Evolution of virulence: triggering host inflammation allows invading pathogens to exclude competitors. Ecology Letters 11. Wiley: 44. doi:10.1111/j.1461-0248.2007.01125.x.

Buckling, A., and P. B. Rainey. 2002. Antagonistic coevolution between a bacterium and a bacteriophage. Proceedings of the Royal Society B Biological Sciences 269. Royal Society: 931. doi:10.1098/rspb.2001.1945.

Cecil, R., E. Ornelas, A. T. Phan, N. O. Medina-Chávez, M. Travisano, and D. R. Yoder-Himes. 2025. Long-term culturing of Pseudomonas aeruginosa in static, minimal nutrient medium results in increased pyocyanin production, reduced biofilm production, and loss of motility. Applied and Environmental Microbiology 91. American Society for Microbiology. doi:10.1128/aem.00975-25.

Chua, S. L., Y. Ding, Y. Liu, Z. Cai, J. Zhou, S. Swarup, D. I. Drautz-Moses, S. C. Schuster, et al. 2016. Reactive oxygen species drive evolution of probiofilm variants in pathogens by modulating cyclic-di-GMP levels. Open Biology 6. Royal Society: 160162. doi:10.1098/rsob.160162.

Coffey, B. M., S. S. Akhand, and G. G. Anderson. 2014. MgtE is a dualfunction protein in Pseudomonas aeruginosa. Microbiology 160. Microbiology Society: 1200. doi:10.1099/mic.0.075275-0.

Coggan, K. A., and M. C. Wolfgang. 2012. Global Regulatory Pathways and Cross-talk Control Pseudomonas aeruginosa Environmental Lifestyle and Virulence Phenotype. Current Issues in Molecular Biology 14. Caister Academic Press: 47. doi:10.21775/cimb.014.047.

Crone, S., M. J. Vives, L. Kvich, A. M. Saunders, M. Malone, M. H. Nicolaisen, E. Martínez-García, C. Rojas-Acosta, et al. 2019. The environmental occurrence of Pseudomonas aeruginosa. Apmis 128. Wiley: 220. doi:10.1111/apm.13010.

Davis, M. R., A. Muszyński, I. V. Lollett, C. L. Pritchett, R. W. Carlson, and J. B. Goldberg. 2013. Identification of the Mutation Responsible for the Temperature-Sensitive Lipopolysaccharide O-Antigen Defect in the Pseudomonas aeruginosa Cystic Fibrosis Isolate 2192. Journal of Bacteriology 195. American Society for Microbiology: 1504. doi:10.1128/jb.01999-12.

Deatherage, D. E., and J. E. Barrick. 2014. Identification of Mutations in Laboratory-Evolved Microbes from Next-Generation Sequencing Data Using breseq. Methods in molecular biology 1151. Springer Science+Business Media: 165. doi:10.1007/978-1-4939-0554-6_12.

Desai, M. M., and D. S. Fisher. 2007. Beneficial Mutation–Selection Balance and the Effect of Linkage on Positive Selection. Genetics 176. Oxford University Press: 1759. doi:10.1534/genetics.106.067678.

Desai, M. M., D. S. Fisher, and A. W. Murray. 2007. The Speed of Evolution and Maintenance of Variation in Asexual Populations. Current Biology 17. Elsevier BV: 385. doi:10.1016/j.cub.2007.01.072.

Diggle, S. P., and M. Whiteley. 2019. Microbe Profile: Pseudomonas aeruginosa: opportunistic pathogen and lab rat. Microbiology 166. Microbiology Society: 30. doi:10.1099/mic.0.000860.

Edgar, R. C. 2022. Muscle5: High-accuracy alignment ensembles enable unbiased assessments of sequence homology and phylogeny. Nature Communications 13. Nature Portfolio: 6968. doi:10.1038/s41467-022-34630-w.

Friman, V., and A. Buckling. 2014. Phages can constrain protist predation-driven attenuation of Pseudomonas aeruginosa virulence in multienemy communities. The ISME Journal 8. Springer Nature: 1820. doi:10.1038/ismej.2014.40.

Fu, Z.-W., F. Ding, B. Zhang, W. Liu, Z.-H. Huang, S. Fan, Y.-R. Feng, Y. Lu, et al. 2024. Hydrogen peroxide sulfenylates and inhibits the photorespiratory enzyme PGLP1 to modulate plant thermotolerance. Plant Communications 5. Elsevier BV: 100852. doi:10.1016/j.xplc.2024.100852.

Gellatly, S. L., B. D. Needham, L. Madera, M. S. Trent, and R. E. W. Hancock. 2012. The Pseudomonas aeruginosa PhoP-PhoQ Two-Component Regulatory System Is Induced upon Interaction with Epithelial Cells and Controls Cytotoxicity and Inflammation. Infection and Immunity 80. American Society for Microbiology: 3122. doi:10.1128/iai.00382-12.

Gerrish, P. J., and R. E. Lenski. 1998. The fate of competing beneficial mutations in an asexual population. PubMed. National Institutes of Health: 127.

Good, B. H., I. M. Rouzine, D. J. Balick, O. Hallatschek, and M. M. Desai. 2012. Distribution of fixed beneficial mutations and the rate of adaptation in asexual populations. Proceedings of the National Academy of Sciences 109. National Academy of Sciences: 4950. doi:10.1073/pnas.1119910109.

Good, B. H., M. J. McDonald, J. E. Barrick, R. E. Lenski, and M. M. Desai. 2017. The dynamics of molecular evolution over 60,000 generations. Nature 551. Nature Portfolio: 45. doi:10.1038/nature24287.

Gouy, M., S. Guindon, and O. Gascuel. 2009. SeaView Version 4: A Multiplatform Graphical User Interface for Sequence Alignment and Phylogenetic Tree Building. Molecular Biology and Evolution 27. Oxford University Press: 221. doi:10.1093/molbev/msp259.

Granato, E. T., C. Ziegenhain, R. L. Marvig, and R. Kümmerli. 2018. Low spatial structure and selection against secreted virulence factors attenuates pathogenicity in Pseudomonas aeruginosa. The ISME Journal 12. Springer Nature: 2907. doi:10.1038/s41396-018-0231-9.

Guindon, S., J.-F. Dufayard, V. Lefort, M. Anisimova, W. Hordijk, and O. Gascuel. 2010. New Algorithms and Methods to Estimate Maximum-Likelihood Phylogenies: Assessing the Performance of PhyML 3.0. Systematic Biology 59. Oxford University Press: 307. doi:10.1093/sysbio/syq010.

Hahn, M. W., E. R. B. Moore, and M. G. Höfle. 2000. Role of Microcolony Formation in the Protistan Grazing Defense of the Aquatic Bacterium Pseudomonas sp. MWH1. PubMed 39. National Institutes of Health: 175. doi:10.1007/s002480000026.

Hickman, J. W., and C. S. Harwood. 2008. Identification of FleQ from Pseudomonas aeruginosa as a c-di-GMP-responsive transcription factor. Molecular Microbiology 69. Wiley: 376. doi:10.1111/j.1365-2958.2008.06281.x.

Hopkins, H. A., C. Lopezguerra, M. Lau, and K. Raymann. 2024. Making a Pathogen? Evaluating the Impact of Protist Predation on the Evolution of Virulence in Serratia marcescens. Genome Biology and Evolution 16. Oxford University Press. doi:10.1093/gbe/evae149.

Hoque, M. M., P. Noorian, G. Espinoza-Vergara, P. M. Cholan, M. Kim, M. H. Rahman, M. Labbate, S. A. Rice, et al. 2021. Adaptation to an amoeba host drives selection of virulence-associated traits in Vibrio cholerae. The ISME Journal 16. Springer Nature: 856. doi:10.1038/s41396-021-01134-2.

Hosseinidoust, Z., N. Tufenkji, and T. G. M. van de Ven. 2013. Predation in Homogeneous and Heterogeneous Phage Environments Affects Virulence Determinants of Pseudomonas aeruginosa. Applied and Environmental Microbiology 79. American Society for Microbiology: 2862. doi:10.1128/aem.03817-12.

Jain, C., L. M. Rodriguez-R, A. M. Phillippy, K. T. Konstantinidis, and S. Aluru. 2018. High throughput ANI analysis of 90K prokaryotic genomes reveals clear species boundaries. Nature Communications 9. Nature Portfolio: 5114. doi:10.1038/s41467-018-07641-9.

Jousset, A., L. Rochat, M. Péchy-Tarr, C. Keel, S. Scheu, and M. Bonkowski. 2009. Predators promote defence of rhizosphere bacterial populations by selective feeding on non-toxic cheaters. The ISME Journal 3. Springer Nature: 666. doi:10.1038/ismej.2009.26.

Kanehisa, M. 2000. KEGG: Kyoto Encyclopedia of Genes and Genomes. Nucleic Acids Research 28. Oxford University Press: 27. doi:10.1093/nar/28.1.27.

Kanehisa, M., Y. Sato, M. Kawashima, M. Furumichi, and M. Tanabe. 2015. KEGG as a reference resource for gene and protein annotation. Nucleic Acids Research 44. Oxford University Press. doi:10.1093/nar/gkv1070.

Kerr, K. G., and A. M. Snelling. 2009. Pseudomonas aeruginosa: a formidable and ever-present adversary. Journal of Hospital Infection 73. Elsevier BV: 338. doi:10.1016/j.jhin.2009.04.020.

Kryazhimskiy, S., D. P. Rice, E. R. Jerison, and M. M. Desai. 2014. Global epistasis makes adaptation predictable despite sequence-level stochasticity. Science 344. American Association for the Advancement of Science: 1519. doi:10.1126/science.1250939.

Leong, W., W. H. Poh, J. G. Williams, C. Lutz, M. M. Hoque, Y. H. Poh, B. Y. K. Yee, C. Chua, et al. 2022. Adaptation to an Amoeba Host Leads to Pseudomonas aeruginosa Isolates with Attenuated Virulence. Applied and Environmental Microbiology 88. American Society for Microbiology. doi:10.1128/aem.02322-21.

Levin, B. R. 1996. The Evolution and Maintenance of Virulence in Microparasites. Emerging infectious diseases 2. Centers for Disease Control and Prevention: 93. doi:10.3201/eid0202.960203.

Marvig, R. L., L. M. Sommer, S. Molin, and H. K. Johansen. 2014. Convergent evolution and adaptation of Pseudomonas aeruginosa within patients with cystic fibrosis. Nature Genetics 47. Nature Portfolio: 57. doi:10.1038/ng.3148.

Matz, C., and S. Kjelleberg. 2005. Off the hook – how bacteria survive protozoan grazing. Trends in Microbiology 13. Elsevier BV: 302. doi:10.1016/j.tim.2005.05.009.

Matz, C., T. Bergfeld, S. A. Rice, and S. Kjelleberg. 2004. Microcolonies, quorum sensing and cytotoxicity determine the survival of Pseudomonas aeruginosa biofilms exposed to protozoan grazing. Environmental Microbiology 6. Wiley: 218. doi:10.1111/j.1462-2920.2004.00556.x.

Matz, C., J. S. Webb, P. J. Schupp, S. Y. Phang, A. Penesyan, S. Egan, P. D. Steinberg, and S. Kjelleberg. 2008a. Marine Biofilm Bacteria Evade Eukaryotic Predation by Targeted Chemical Defense. PLoS ONE 3. Public Library of Science. doi:10.1371/journal.pone.0002744.

Matz, C., A. M. Moreno, M. Alhede, M. Manefield, A. R. Hauser, M. Givskov, and S. Kjelleberg. 2008b. Pseudomonas aeruginosa uses type III secretion system to kill biofilm-associated amoebae. The ISME Journal 2. Springer Nature: 843. doi:10.1038/ismej.2008.47.

Mikonranta, L., V. Friman, and J. Laakso. 2012. Life History Trade-Offs and Relaxed Selection Can Decrease Bacterial Virulence in Environmental Reservoirs. PLoS ONE 7. Public Library of Science. doi:10.1371/journal.pone.0043801.

Molina, K. J. A., S. J. Gutiérrez, N. Benítez-Campo, and A. Correa. 2024. Genomic Differences Associated with Resistance and Virulence in Pseudomonas aeruginosa Isolates from Clinical and Environmental Sites. Microorganisms 12. Multidisciplinary Digital Publishing Institute: 1116. doi:10.3390/microorganisms12061116.

Nair, R. R., M. Vasse, S. Wielgoss, L. Sun, Y.-T. N. Yu, and G. J. Velicer. 2019. Bacterial predator-prey coevolution accelerates genome evolution and selects on virulence-associated prey defences. Nature Communications 10. Nature Portfolio. doi:10.1038/s41467-019-12140-6.

Rainey, P. B., and M. Travisano. 1998. Adaptive radiation in a heterogeneous environment. Nature 394. Nature Portfolio: 69. doi:10.1038/27900.

Rambaut, A. 2018. FigTree v1.4.4.Institute of Evolutionary Biology, University of Edinburgh, http://tree.bio.ed.ac.uk/software/figtree/.

Ravichandran, A., M. Ramachandran, T. Suriyanarayanan, J. C. C. Wong, and S. Swarup. 2015. Global Regulator MorA Affects Virulence-Associated Protease Secretion in Pseudomonas aeruginosa PAO1. PLoS ONE 10. Public Library of Science. doi:10.1371/journal.pone.0123805.

Saha, S., S. Sureshkumar, V. Sharma, and S. Pande. 2025. Coevolutionary history of predation constrains the evolvability of antibiotic resistance in prey bacteria. npj Antimicrobials and Resistance 3: 49. doi:10.1038/s44259-025-00111-5.

Schwengers, O., L. Jelonek, M. A. Dieckmann, S. Beyvers, J. Blom, and A. Goesmann. 2021. Bakta: rapid and standardized annotation of bacterial genomes via alignment-free sequence identification. Microbial Genomics 7. Microbiology Society. doi:10.1099/mgen.0.000685.

Silby, M. W., C. Winstanley, S. A. C. Godfrey, S. B. Levy, and R. W. Jackson. 2011. Pseudomonasgenomes: diverse and adaptable. FEMS Microbiology Reviews 35. Oxford University Press: 652. doi:10.1111/j.1574-6976.2011.00269.x.

Stover, C. K., X. Q. Pham, A. L. Erwin, S. Mizoguchi, P. Warrener, M. J. Hickey, F. S. L. Brinkman, W. O. Hufnagle, et al. 2000. Complete genome sequence of Pseudomonas aeruginosa PAO1, an opportunistic pathogen. Nature 406. Nature Portfolio: 959. doi:10.1038/35023079.

Tonkin-Hill, G., N. MacAlasdair, C. Ruis, A. Weimann, G. Horesh, J. A. Lees, R. A. Gladstone, S. W. Lo, et al. 2020. Producing polished prokaryotic pangenomes with the Panaroo pipeline. Genome biology 21. BioMed Central: 180. doi:10.1186/s13059-020-02090-4.

Wahl, L. M., and P. J. Gerrish. 2001. THE PROBABILITY THAT BENEFICIAL MUTATIONS ARE LOST IN POPULATIONS WITH PERIODIC BOTTLENECKS. Evolution 55. Oxford University Press: 2606. doi:10.1111/j.0014-3820.2001.tb00772.x.

Williams, H. T. P. 2013. Phage-induced diversification improves host evolvability. BMC Evolutionary Biology 13. Springer Science+Business Media: 17. doi:10.1186/1471-2148-13-17.

Wu, M., and X. Li. 2014. Klebsiella pneumoniae and Pseudomonas aeruginosa. Elsevier eBooks. Elsevier BV: 1547. doi:10.1016/b978-0-12-397169-2.00087-1.

