## Supplemental Files and Figures for "Genomic and phenotypic diversification of *Pseudomonas aeruginosa* during sustained exposure to a ciliate predator"

**Supplemental Material**

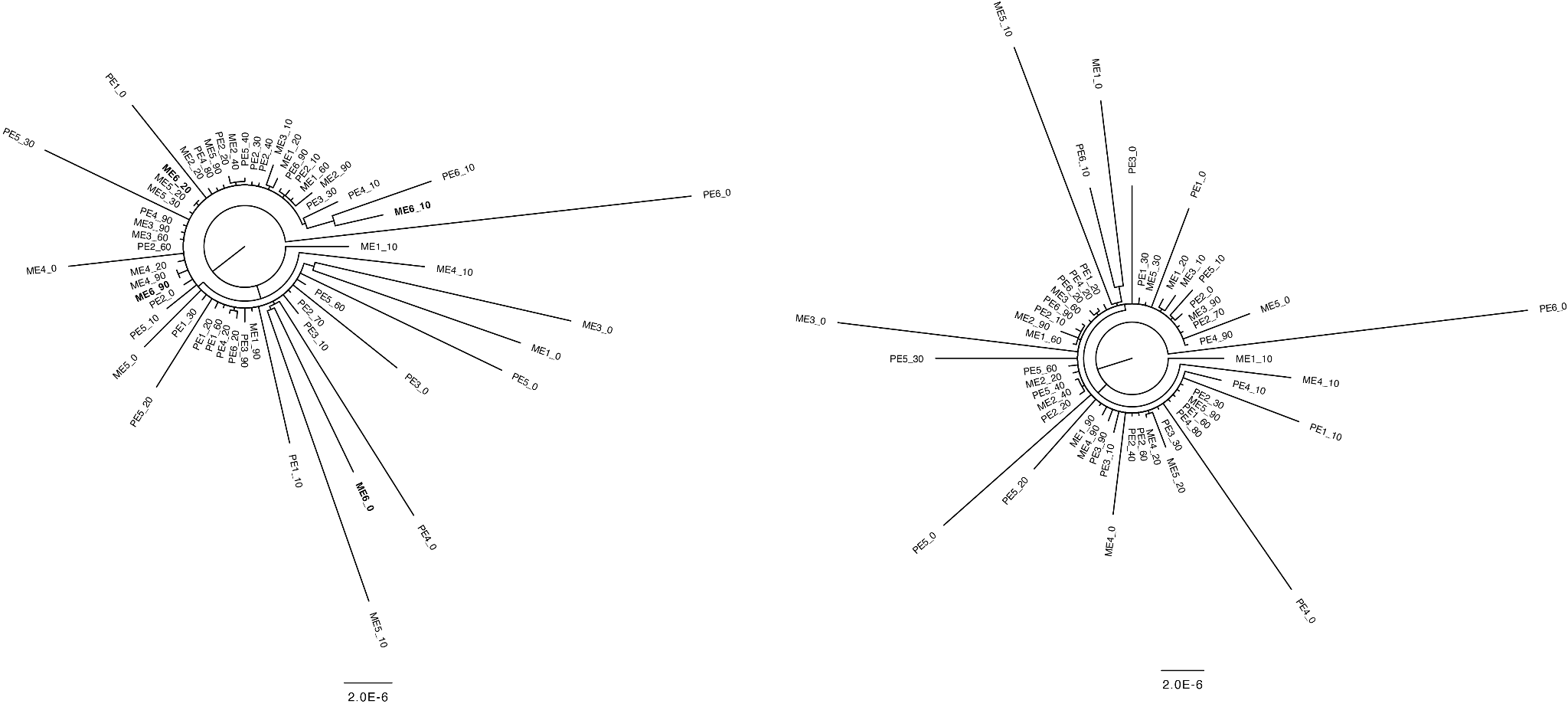

**Figure S1. Phylogenetic analysis of pseudoclone genomes and detection of cross-contamination among evolved populations.** Phylogenetic relationships among evolved populations were inferred using pseudoclones constructed from allele frequency bins (see Methods). Briefly, within each population, variants were grouped into frequency bins (<10%, 10–20%, 20–30%, …, >90%), and bin-specific mutations were used to generate pseudoclone genomes, which were then aligned using MUSCLE v5 and analysed phylogenetically using PhyML v3.0 under a GTR model. Trees were visualized in FigTree v1.4.4. The left panel shows the phylogeny including all pseudoclones, while the right panel shows the same analysis with ME6 removed due to evidence of cross-contamination.

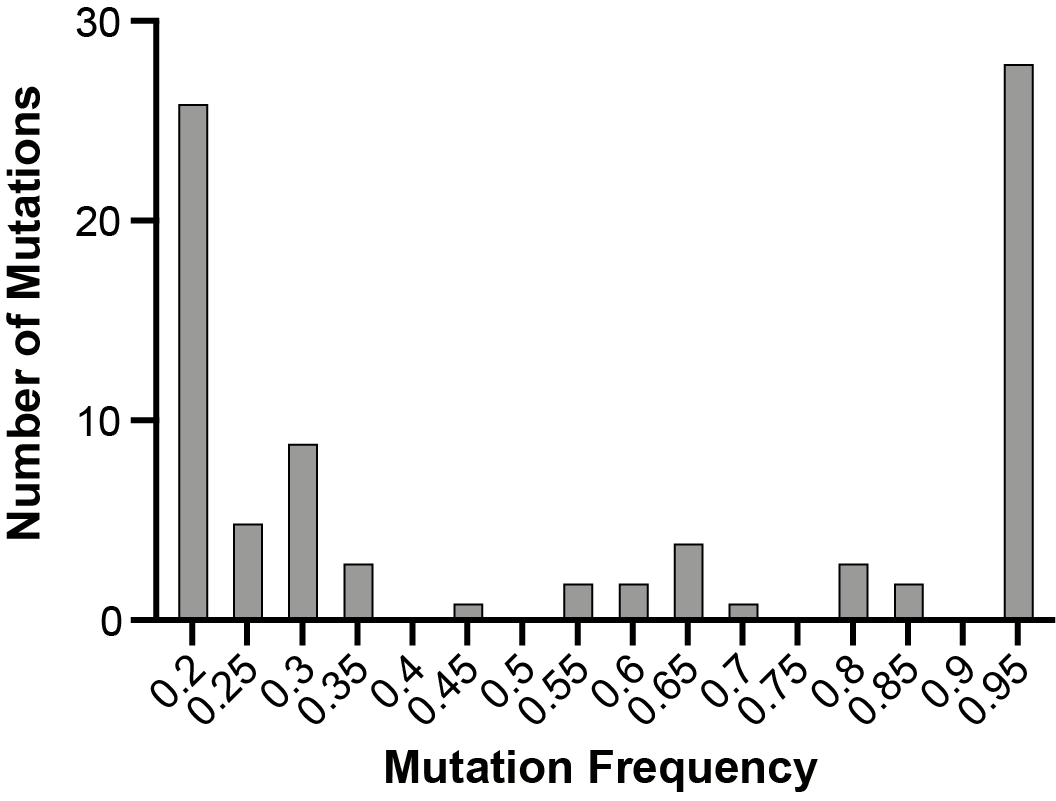

**Figure S2. Distribution of mutation frequencies (≥0.20).** Number of mutations detected across all evolved populations with frequencies ≥0.20, binned into 0.05 intervals. Each bar represents the lower bound of the corresponding frequency bin (e.g., 0.20 = 0.20–0.24; 0.95 = 0.95–1.00).

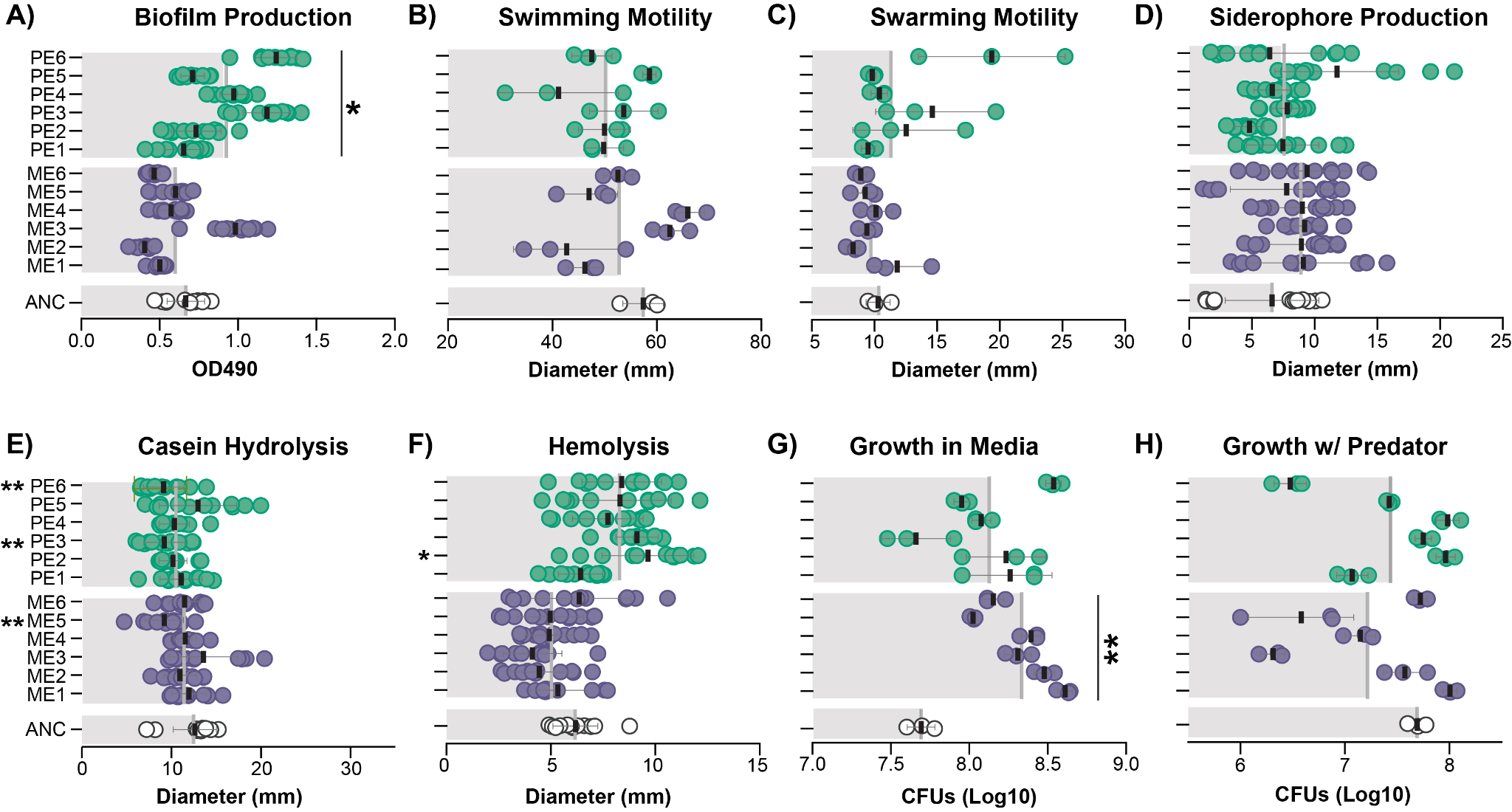

**Figure S3. Phenotypic characteristics of evolved isolates.** Assays performed were: (A) biofilm production, (B) swimming motility, (C) swarming motility, (D) siderophore production, (E) casein hydrolysis, (F) hemolytic activity, (G) growth in media after 24 h based on CFU counts, and (H) growth in coculture with *T. thermophila* after 24 h based on CFU counts. Statistical significance was determined using a Kruskal–Wallis test, followed by Dunn’s multiple comparisons test. For all analyses, the mean rank of each group was compared to that of the ancestral strain. Comparisons were performed for each evolved isolate individually (asterisks on the left) and for the mean of all ME and PE isolates (asterisks on the right) relative to the mean of the ancestor. *, P < 0.05; **, P < 0.01.

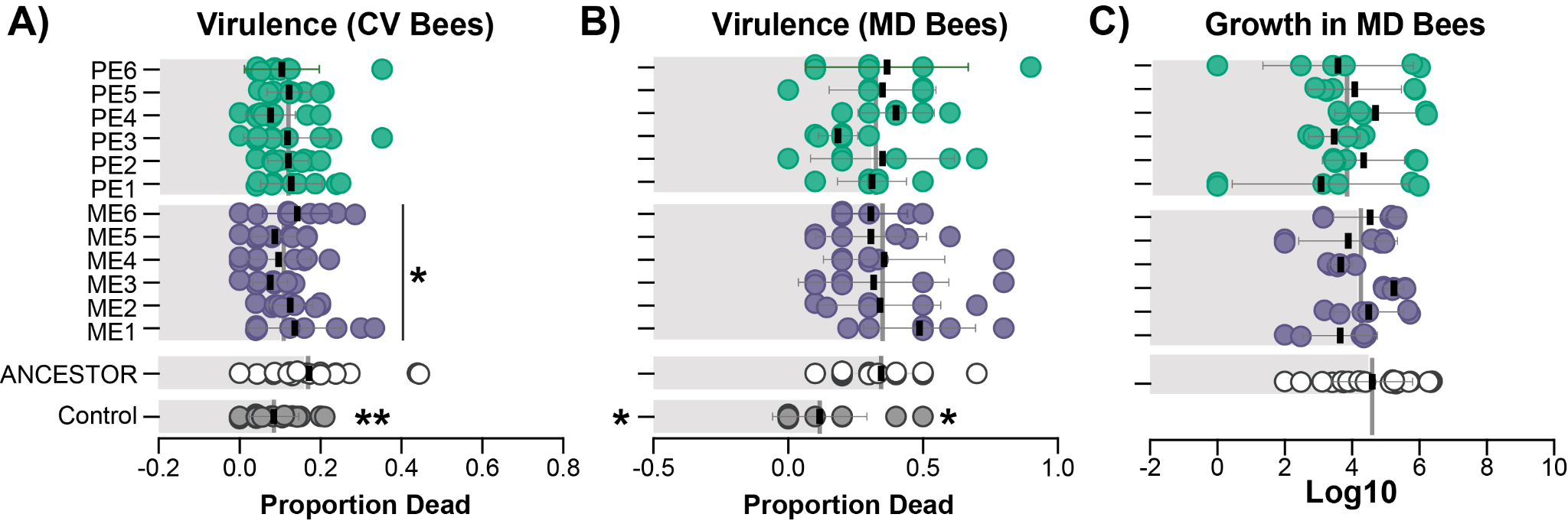

**Figure S4. Virulence and growth of evolved isolates in honey bees.** Proportion of dead (A) conventional (CV) honey bees and (B) microbiota-depleted (MD) honey bees after 5 days of exposure to the ancestral strain or individual PE and ME isolates. (C) Growth of the ancestor, PE, and ME isolates in microbiota-depleted honey bees based on colony-forming unit (CFU) counts; CFU values are shown as log₁₀-transformed counts. In panels A and B, each data point represents a cup cage containing 20 bees for CV trials and 10 bees for MD trials (biological replicates with technical replication). In panel C, each data point represents an individual bee. Statistical significance was tested both between each individual PE and ME isolate and the ancestral strain, and between the mean of PE and ME isolates and the ancestral strain. Statistical significance was determined using a Kruskal–Wallis test followed by Dunn’s multiple comparisons test (*, P < 0.05; **, P < 0.01).

| **Sample** | **Line** | **Average Coverage** | **% Aligned** |
| --- | --- | --- | --- |
| Isolate | PE-1 | 31.90703 | 99.45% |
| Isolate | PE-2 | 44.70393 | 99.37% |
| Isolate | PE-3 | 32.88026 | 99.42% |
| Isolate | PE-4 | 28.97971 | 98.98% |
| Isolate | PE-5 | 39.65268 | 98.99% |
| Isolate | PE-6 | 24.55709 | 98.96% |
| Isolate | ME-1 | 19.68769 | 99.06% |
| Isolate | ME-2 | 47.9632 | 99.19% |
| Isolate | ME-3 | 93.0591 | 99.15% |
| Isolate | ME-4 | 35.00678 | 99.02% |
| Isolate | ME-5 | 29.0038 | 98.59% |
| Isolate | ME-6 | 55.80903 | 99.05% |
| Population | PE-1 | 38.33171 | 99.15% |
| Population | PE-2 | 282.7614 | 99.21% |
| Population | PE-3 | 49.2864 | 98.80% |
| Population | PE-4 | 66.66734 | 98.29% |
| Population | PE-5 | 244.5668 | 98.99% |
| Population | PE-6 | 91.32875 | 97.91% |
| Population | ME-1 | 81.95025 | 98.53% |
| Population | ME-2 | 39.79088 | 98.71% |
| Population | ME-3 | 89.21901 | 98.66% |
| Population | ME-4 | 38.41933 | 98.57% |
| Population | ME-5 | 30.37836 | 97.94% |
| Population | ME-6 | 71.459 | 98.28% |

**Table S1. Average genome coverage and percent of reads aligned for the evolved populations and isolates.**

| **Line** | **NumMut** | **Generations** | **Mutations/generation** |
| --- | --- | --- | --- |
| ME1_60 | 49 | 276 | 0.1775 |
| ME2_60 | 10 | 276 | 0.0362 |
| ME3_60 | 54 | 276 | 0.1957 |
| ME4_60 | 60 | 276 | 0.2174 |
| ME5_60 | 84 | 276 | 0.3043 |
| PE1_60 | 84 | 276 | 0.3043 |
| PE2_60 | 12 | 276 | 0.0435 |
| PE3_60 | 51 | 276 | 0.1848 |
| PE4_60 | 92 | 276 | 0.3333 |
| PE5_60 | 68 | 276 | 0.2464 |
| PE6_60 | 98 | 276 | 0.3551 |

**Table S2. Estimated number of mutations per generation.**

**Dataset S1. Mutations identified in evolved populations.** List of mutations detected across all evolved populations at a frequency greater than 5%.

**Dataset S2. High frequency mutations in evolved populations.** List of mutations detected at >=20% in at least one population.

**Dataset S3. Mutations identified in evolved isolates.** List of mutations detected across all evolved isolates.
